# Single-step purification of functionalized protein nanostructures using multimodal chromatography

**DOI:** 10.1101/2022.08.21.504681

**Authors:** Daniel L. Winter, Hélène Lebhar, Joshua B. McCluskey, Dominic J. Glover

## Abstract

Protein nanostructures produced through the self-assembly of individual subunits are attractive scaffolds to attach and position functional molecules for applications in biomaterials, metabolic engineering, tissue engineering, and a plethora of nanomaterials. However, the assembly of multicomponent protein nanomaterials is generally a laborious process that requires each protein component to be separately expressed and purified prior to assembly. Moreover, excess components not incorporated into the final assembly must be removed from the solution and thereby necessitate additional processing steps. Here, we developed an efficient approach to purify functionalized protein filament assemblies directly from bacterial lysates in a single step through a type of multimodal chromatography that combines size-exclusion, hydrophilic interaction, and ion exchange to separate recombinant protein assemblies from excess free subunits and bacterial proteins. In this approach, the ultrastable filamentous protein gamma-prefoldin was employed as a material scaffold that can be functionalized with a variety of protein domains through SpyTag/SpyCatcher conjugation chemistry. The purification of recombinant gamma-prefoldin filaments from bacterial lysates using multimodal chromatography was optimized across a wide range of salt concentrations and pH. Subsequently, functionalized protein assemblies were purified from bacterial lysates using multimodal chromatography in a single step and shown to befree of unincorporated subunits. The assembly and purification of protein nanostructures with varying amounts of functionalization was confirmed using polyacrylamide gel electrophoresis, Förster resonance energy transfer, and transmission electron microscopy. We envision that the use of multimodal chromatography will increase the throughput of protein nanostructure prototyping as well as enable the upscaling of the bioproduction of protein nanodevices.

## Introduction

The self-assembly of proteins can be harnessed to produce intricate nanostructures that can be functionalized with additional protein domains to create nanodevices for a variety of applications. Functionalized protein nanostructures are generally composed of many different protein subunits, with some subunits assembling as an architectural scaffold, while other subunits serve as functional components to produce chemical activities. The functional protein subunits are attached to the scaffold, whereby their proximity and alignment results in emergent properties of the whole assembly. The alignment of functional domains on protein nanostructures includes enzymes for multistep biocatalysis pathways (1–3), cell signaling domains for tissue engineering (4), and metalloproteins on filaments to produce electronically conductive nanowires (5), However, the fabrication of multicomponent protein complexes is laborious, as the different components must be recombinantly expressed and purified separately before assembly of the functional nanostructure can occur (6).

It would be advantageous to the field of nanobiotechnology if the fabrication steps required to produce a nanostructure could be reduced and the process standardized using scalable manufacturing methodology. In the case of a homomeric protein nanostructure such as a symmetrical cage, only one protein component needs to be expressed and assembled. In this simple scenario, protein production is typically performed by recombinant expression in a microbial host, whereupon the protein monomer self-assembles *in vivo* into a pre-defined structure (7). However, more useful assemblies can be obtained by combining different protein components, including functional domains into structurally complex heteromeric nanostructures (8). Our capacity to engineer intricate self-assembling protein complexes has dramatically increased with the design of sequence-specific binding domains (9–11), and orthogonal reactive interfaces that form covalent attachments between proteins (12, 13). However, the production of heteromeric protein assemblies requires laborious procedures to separately express and purify each individual protein component, followed by nanostructure assembly and removal of unincorporated protein subunits. We thus sought to develop an assembly and purification strategy to purify heteromeric protein assemblies from bacterial lysates, which is also compatible with a broad range of pH and salt conditions and does not rely on the incorporation of chromatography affinity tags.

Capto Core 700 (CC700) is a multimodal chromatography (MMC) resin combining size-exclusion, ion exchange, and hydrophobic interaction chromatography to enable purification of large protein assemblies. The resin is composed of beads whose surface is inactive and cannot bind to biomolecules.

The pores of the CC700 beads have a large molecular weight cut-off (700 kDa), whereby assemblies larger than the pore size are excluded from entering the beads and are instead recovered in the chromatography flow-through. Smaller molecular assemblies, however, enter the pores and are retained inside the beads through interaction with the internal bead surface that has a mixture of hydrophilic and hydrophobic groups for molecule binding.

Previously, CC700 has been used to purify viruses from mammalian cell lysates (14), and recombinantly expressed viral-like particles (15, 16) and encapsulin nanocompartments (17). However, we hypothesize that CC700 technology can be expanded to enable single-step assembly and purification of a functionalized heteromeric protein nanostructure. Here, we verified the robustness of our strategy by systematically comparing the purification yields of the protein gamma-prefoldin (γPFD), which self-assembles into filaments in the megadalton scale, at several salt concentrations and pH values. Subsequently, we demonstrated the filaments could be functionalized and purified directly from bacterial lysates. An engineered gamma-prefoldin fused to a SpyTag domain and three functional proteins fused to a SpyCatcher domain were recombinantly produced in separate *Escherichia coli* strains. Combining the four bacterial lysates triggered nanostructure assembly and functionalization (Figure 1a) and enabled the full assembly to be purified using MMC, free of excess subunits that were not incorporated into the complex (Figure 1b). To further improve purity, sample processing steps were tested both upstream and downstream of MMC, including tangential flow filtration (TFF) and Triton X-114 phase separation, which increased resin capacity or removed any remaining lipids and protein contaminants. The capacity to assemble complex assemblies and their subsequent purification using a rapid, standardized procedure will facilitate the production and upscaling of engineered protein nanostructures.

**Figure 1.**
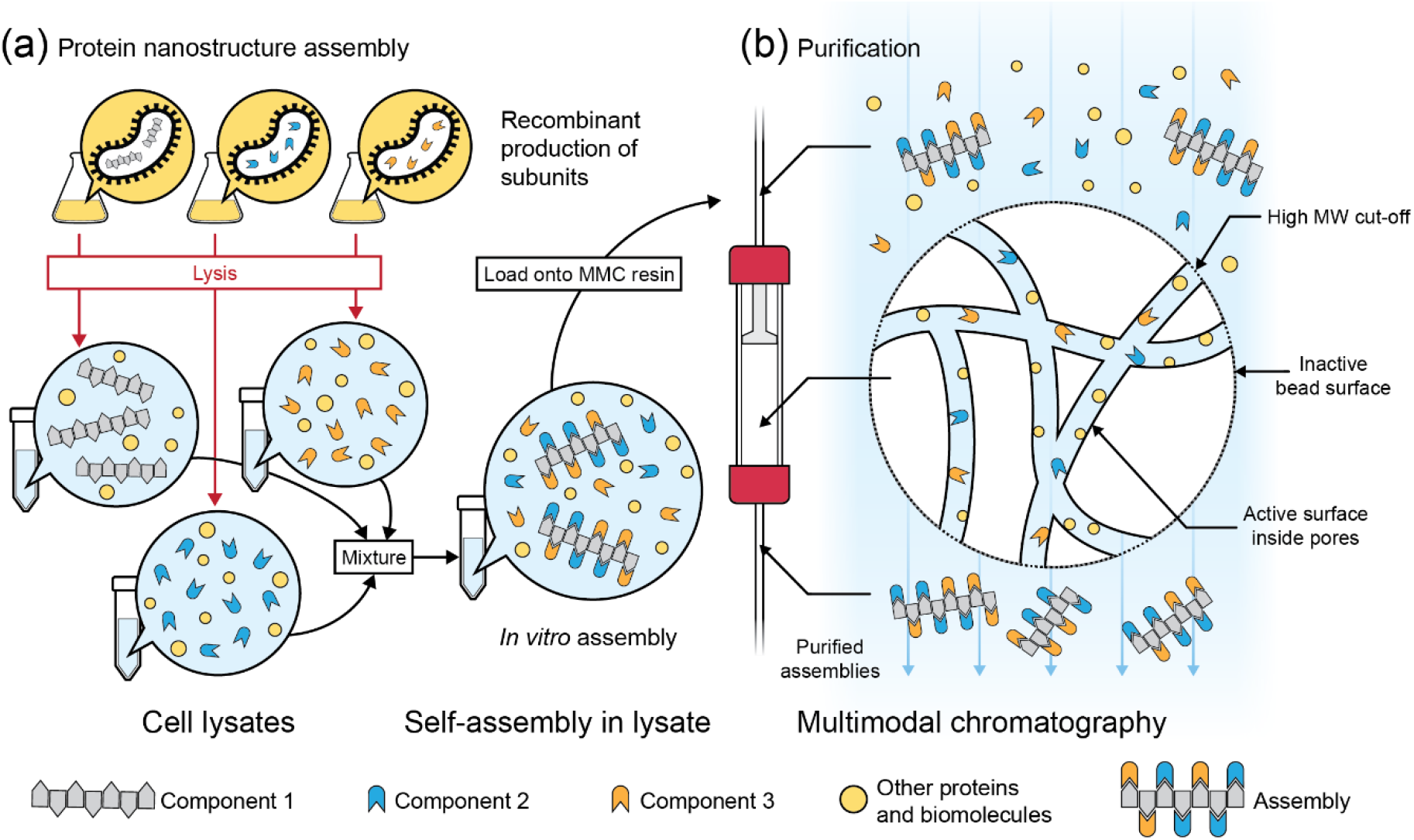
Recombinant production of protein nanostructures and their purification by multimodal chromatography. **(a)** The components of a protein assembly can be recombinantly produced in separate microbial strains. Following lysis of each bacterial culture, the lysates are mixed, wherein the components self-assemble into nanostructures. **(b)** Following assembly, the protein nanostructures can be purified from the rest of the lysate, including excess free components, using a multimodal chromatography (MMC) resin, Capto Core 700 (CC700). In CC700 MMC, only molecules that are smaller than the resin pores can enter the beads, where they bind to the active surface inside the beads. Because the assembled nanostructures are too large to enter the pores, and because the shell of the beads is inactive, the nanostructures do not bind to the resin and are recovered in the flow-through.

## Materials and methods

### Preparation of bacterial expression strains

Plasmids for bacterial expression of wild type γPFD, γPFD-SpyTag-6×His, SpyCacther-6×His, mCerulean3-SpyCatcher-6×His, and mVenus-SpyCatcher-6×His were prepared using Gibson assembly with pET-19b (Novagen) as the plasmid vector. A linear pET-19b backbone was produced by polymerase chain reaction (PCR), and the insert DNA fragments were prepared by either PCR or commercially synthesized by Integrated DNA Technologies. Assembled DNA plasmids were transferred into competent T7 Express *E. coli* cells (New England Biolabs) and colonies selected on lysogeny broth (LB) agar plates supplemented with 50 μg/mL of ampicillin. The fidelities of expression plasmids were verified by Sanger sequencing at the Ramaciotti Centre for Genomics at the University of New South Wales.

### Expression of individual protein components and preparation of lysates

Overnight cultures were prepared for each strain by inoculating 5 mL LB medium supplemented with 50 μg/mL of ampicillin with a single colony. The next day, flasks containing 100 mL of LB medium with ampicillin were inoculated with 1 mL of an overnight culture and grown at 37 °C with shaking until an optical density of 0.6 at A600. Subsequently, protein expression was induced for 15 h at 20 °C with the addition of isopropyl β-d-1-thiogalactopyranoside (IPTG) to a final concentration of 1 mM. Cells were harvested by centrifugation and lysis was performed by sonication in phosphate-buffered saline (50 mM sodium phosphate, 100 mM sodium chloride, pH 8.0). Cell debris were removed by centrifugation at 20,000 *g* followed by filtration using 0.22 μm Millex syringe filters (Merck). Cleared lysates were used for subsequent protein assembly experiments.

A similar procedure was performed for large-scale expression of wild type γPFD, except a total of 15 L of LB medium were inoculated for protein expression and lysis was performed in phosphate-buffered saline using a French press, for a total of 2 L of lysate. Bovine deoxyribonuclease I (Sigma-Aldrich) was added to a final concentration of 0.1 mg/mL to degrade genomic and plasmid DNA and reduce the viscosity of the lysate. The lysate was clarified by centrifugation at 10,000 *g* and subjected to tangential flow filtration (TFF) as described below. The TFF-processed lysate was split into volumes of 150 mL, which were either stored at 4 °C or buffer-exchanged using SnakeSkin dialysis tubing (Thermo Fisher) with a 10 kDa cut-off prior to storage. The buffers used for buffer exchange are indicated in Table 1.

**Table 1.**
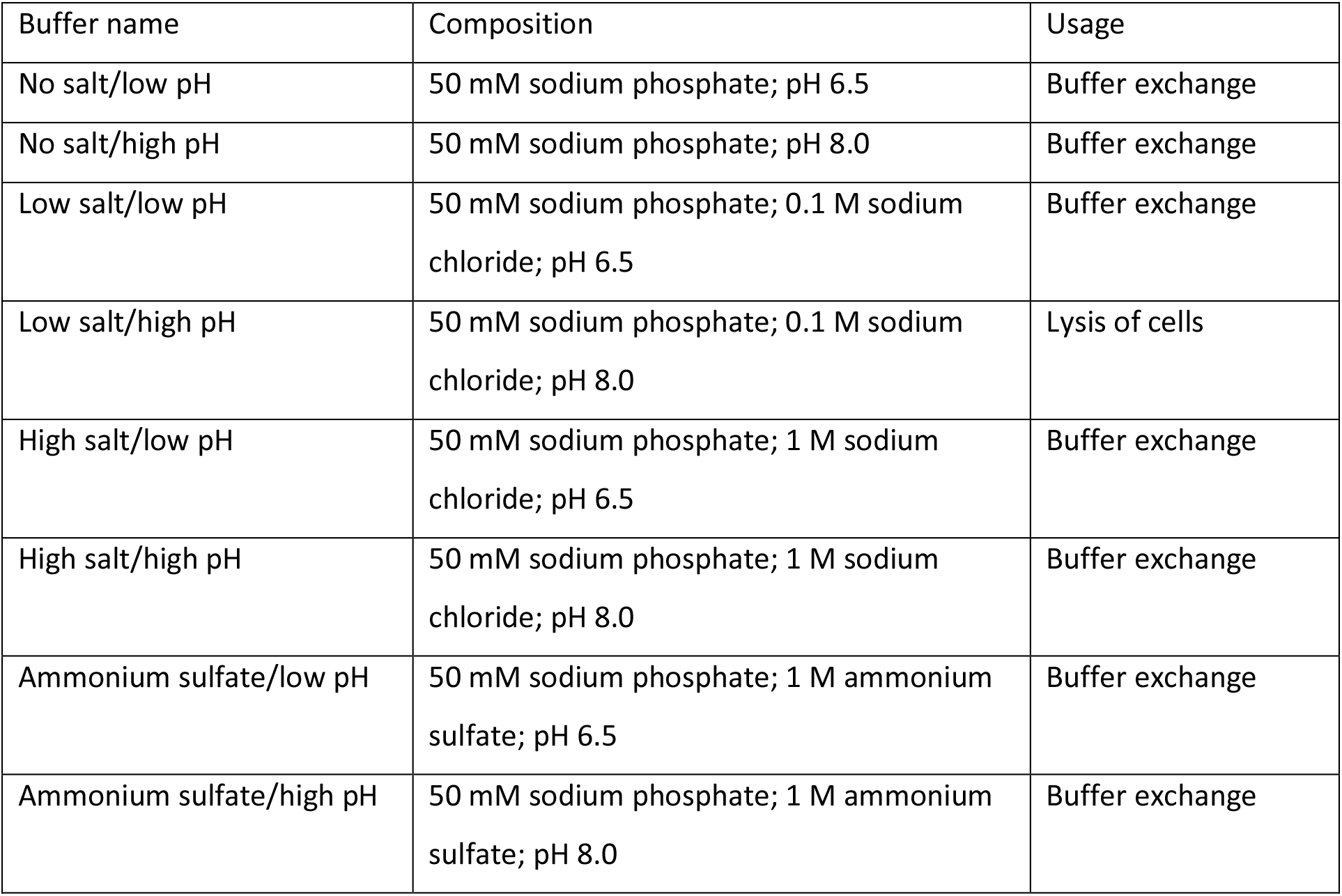
Buffered solvents used for CC700 purification.

| Buffer name | Composition | Usage |
| --- | --- | --- |
| No salt/low pH | 50 mM sodium phosphate; pH 6.5 | Buffer exchange |
| No salt/high pH | 50 mM sodium phosphate; pH 8.0 | Buffer exchange |
| Low salt/low pH | 50 mM sodium phosphate; 0.1 M sodium chloride; pH 6.5 | Buffer exchange |
| Low salt/high pH | 50 mM sodium phosphate; 0.1 M sodium chloride; pH 8.0 | Lysis of cells |
| High salt/low pH | 50 mM sodium phosphate; 1 M sodium chloride; pH 6.5 | Buffer exchange |
| High salt/high pH | 50 mM sodium phosphate; 1 M sodium chloride; pH 8.0 | Buffer exchange |
| Ammonium sulfate/low pH | 50 mM sodium phosphate; 1 M ammonium sulfate; pH 6.5 | Buffer exchange |
| Ammonium sulfate/high pH | 50 mM sodium phosphate; 1 M ammonium sulfate; pH 8.0 | Buffer exchange |

### Tangential flow filtration

Tangential flow filtration was performed using a Labscale TFF System (Merck) and a Pellicon XL50 with Ultracel 1000 kDa membrane (Merck) using a feed flow rate of 20 mL/min and a transmembrane pressure maintained at 3-5 psi. A constant volume diafiltration was performed, where the retentate volume was held constant at 250 mL and the diafiltration buffer was added to the retentate at the same flow rate as the permeate. The experiment was stopped once 4 volumes of buffer were diafiltered with the total permeate volume reaching 1 L. Sampling of retentate and permeate was undertaken every 50 mL of permeate produced using sampling valves located on the retentate and permeate port of the diafiltration cassette.

### Purification by multimodal chromatography

For the testing of different solvents, MMC was performed with 1 mL HiTrap columns loaded with Capto Core 700 (Cytiva) and an ÄKTA pure 25 FPLC system (Cytiva) to automate the process. Each column was equilibrated with 10 column volumes (CV) of one of eight buffers (see Table 1) followed by the application of 20 CV of bacterial lysate (previously dialyzed in the same solvent, see above) containing recombinant γPFD. The flow-through was collected in 1 mL volume fractions. After each run, the columns were cleaned with 10 CV of distilled water, 10 CV of a cleaning-in-place solvent (1 M sodium hydroxide in 30% (v/v) isopropanol), 10 CV of distilled water, and re-equilibrated with one of the eight buffers being tested (Table 1). All steps were performed at 0.5 mL/min flow rate. In all other experiments, MMC was performed by gravity flow using Poly-Prep Chromatography Columns (Bio-Rad) loaded with up to 2 mL of Capto Core 700 resin. The procedure was essentially the same, except all steps were performed manually.

### Fluorescence measurements

Fluorescence and Förster resonance energy transfer (FRET) measurements were performed with a CLARIOstar plate reader (BMG Labtech). For the measurement of fluorescence emission spectra, the excitation wavelength was set 433 nm (10 nm width) and the fluorescence emission was recorded from 455 nm to 600 nm wavelengths (10 nm width). For the measurement of FRET emission, three fluorescence values were recorded, with the following excitation/emission wavelengths: 433/476 nm (mCerulean3), 515/527 nm (mVenus), and 433/527 nm (FRET). The method described by Song *et al.* (18) was used to subtract background fluorescence resulting from the direct activation of mCerulean3 and mVenus and obtain the value of the FRET emission.

### Transmission electron microscopy

Samples were diluted to approximately 20 μg/mL and deposited on carbon type-B, 200-mesh copper grids (Ted Pella Inc.). Grids had previously been treated with an easiGlow glow discharge unit (PELCO). After a 5 min incubation at room temperature, the grids were washed with distilled water and stained with 2% aqueous uranyl acetate (BDH Chemicals) for 7 min. Excess uranyl acetate was absorbed with filter paper (Whatman), and the grids were left to dry before visualization under a Tecnai G2 20 TEM (FEI).

## Results and discussion

### CC700 enables robust purification of recombinant protein filaments from bacterial lysates

The combined assembly and purification of functional protein nanostructures from cell lysates will require solvent conditions that are compatible for each process. Thus, we performed a series of experiments to determine whether a commercially available MMC resin, Capto Core 700 (CC700), could efficiently purify protein nanostructures in varying pH and salt conditions and identify the optimal conditions for the purification of large protein assemblies. Specifically, we chose salt concentrations ranging from 0 to 1 M sodium chloride or 1 M ammonium sulfate, and pH values of either 6.5 or 8.0. The choice of salt, salt concentration, and pH should modulate the binding of proteins to the ion exchange and hydrophobic sites of CC700 resin (19) as well as the stability of large protein assemblies, and thereby affects the purity of the recovered protein nanostructures.

The archaeal molecular chaperone γPFD, which self-assembles into micrometer-long filaments and has previously been decorated with a variety of functional molecules (1, 5, 20, 21) was chosen as a protein nanostructure. Filaments of γPFD can be recombinantly produced (Figure 2a) and are stable across a broad range of environmental conditions, which enabled us to compare the purification yield of the filaments using CC700 in the chosen range of salt and pH conditions. Filaments of γPFD were recombinantly expressed in *E. coli* and the cell pellets were lysed in a low salt, phosphate-buffered solution. An aliquot of the lysate was subjected to TFF with a large molecular weight cut-off (1,000 kDa) to compare the performance of TFF and CC700 for the removal of bacterial proteins from the lysate. TFF did not efficiently remove bacterial proteins from the γPFD-containing lysate, however nucleic acid content in the lysate was drastically reduced (Figure 2b and Figure S1). Presumably, because the lysate was treated with the endonuclease DNase I, the genomic and plasmid DNA were cleaved into small oligonucleotides that passed through the TFF filter. The TFF-processed lysate was then divided and dialyzed into buffer solutions that varied in salt content and pH, totaling nine different loading conditions for MMC purification (Figure 2c). The individual samples were loaded onto pre-packed 1 mL CC700 columns, with similar protein content and concentration (Figure S2). Chromatograms of MMC purifications indicated that protein content in the flow-through quickly increased after a few column volumes passed through the CC700 column, suggesting saturation of the resin, at which point the resin would cease to retain small proteins and the flow-through would not contain pure γPFD. We analyzed the early fractions from two MMC purification conditions (with and without TFF processing) by SDS-PAGE and confirmed that only the first few flow-through fractions contained purified γPFD and that TFF processing of the lysate improved purity (Figure S3). Two early fractions from each MMC purification condition were analyzed by SDS-PAGE to compare the purity of γPFD, which confirmed that the sample that was not subjected to TFF and thus contained higher nucleic acid concentration showed lesser purity compared to TFF-processed loads (Figure 2d, Figure S3, Figure S4). Presumably, oligonucleotides and other small biomolecules from the non-TFF-processed lysate were retained in the CC700 resin, leading to earlier saturation of the column binding capacity and cessation of effective purification.

**Figure 2.**
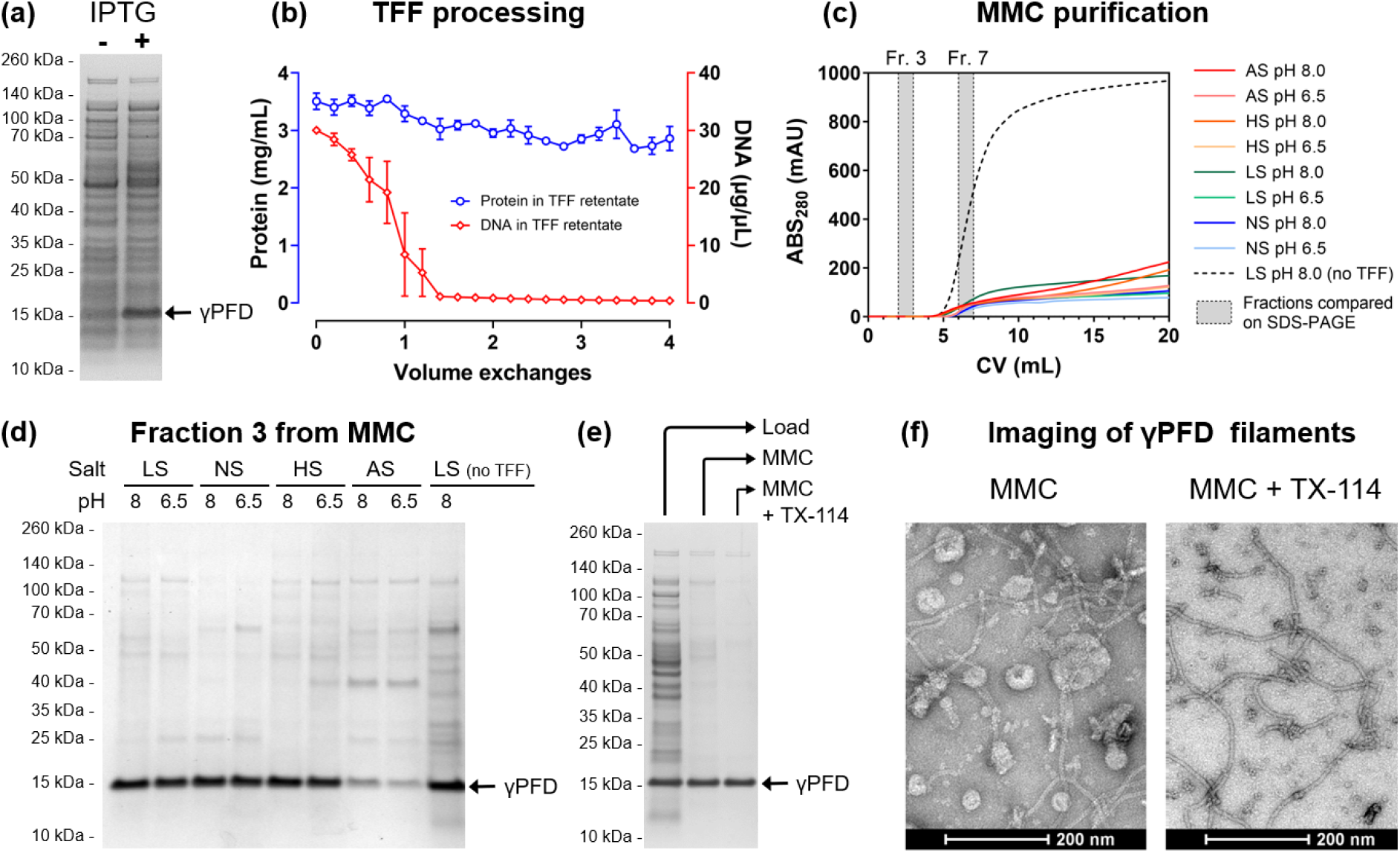
CC700 resin enables consistent protein assembly purification across a range of pH and salt conditions. **(a)** Recombinant expression of γPFD in E. coli following inducement with IPTG as shown by SDS-PAGE. **(b)** TFF with a 1 MDa filter does not remove contaminant proteins from a bacterial lysate but is effective in removing nucleic acids. **(c)** UV chromatograms of MMC purifications showing comparable performance in a range of buffered solvents. Pre-processing with TFF increases resin binding capacity. **(d)** Comparison of an early fraction from MMC performed with different buffered solvents confirms that MMC performs comparably in all solvents except for ammonium sulfate. Omitting TFF pre-processing results in lesser purity. **(e)** Large protein assemblies purified using MMC can be further purified with a TX-114 phase separation procedure. **(f)** Transmission electron microscopy reveals contamination, possibly lipidic in nature, without TX-114 post-processing. LS: low salt (100 mM NaCl); NS: no salt; HS: high salt (1 M NaCl); AS: ammonium sulfate (1 M); TX-114: Triton X-114.

Additional evidence of increased CC700 capacity following lysate processing by TFF are the elution chromatograms for each tested condition (Figure 2c and Figure S5), which show a much steeper increase in UV absorbance at 280 nm across fractions for the sample that was not pre-processed using TFF compared to the TFF-processed conditions. Thus, TFF served to increase the binding capacity of CC700 resin for bacterial proteins by removing biomolecules (predominantly nucleic acids), which is useful for the scale up of MMC purifications. MMC buffer conditions that used 1 M ammonium sulfate instead of sodium chloride resulted in less recovery of γPFD (Figure 2d). The high ionic strength of ammonium sulfate compared to sodium chloride may lead to some precipitation of γPFD or interfere with filament formation resulting in retention of γPFD monomers in the resin, explaining the lower recovery.

Nonetheless, all purification conditions produced a trace amount of background protein contamination (Figure 2d). We reasoned that protein contamination may result from proteins bound to bacterial lipid membrane fragments that are known to form vesicles cell lysates. The nonionic detergent Triton-X114 is commonly used to remove endotoxins (which are lipopolysaccharides that can aggregate into vesicles) from bacterial lysates and has been previously used as a downstream polishing step following MMC for the preparation of virus-like particles (15). We thus attempted to improve the purity of γPFD following MMC by phase-separating lipids from proteins using Triton X-114 (22). The simple procedure was shown to remove background contamination and improve the purity of γPFD (Figure 2e). In addition, transmission electron microscopy (TEM) showed that non-protein particle contaminants, possibly lipidic in nature, were removed from MMC-purified γPFD following Triton X-114 phase separation (Figure 2f). Altogether, CC700 resin performed similarly well for most salt and pH conditions and should thus be compatible with most types of functionalized protein assemblies. Straightforward bacterial lysate pre- and post-processing using TFF and Triton-X114 phase separation, respectively, can be used to improve the capacity of MMC resin and the purification of large protein assemblies.

### Strategy to attach and interspace functional proteins to γPFD filaments

Single-step assembly and purification of functionalized protein nanostructures from bacterial lysates would be a useful methodology for the field of nanobiotechnology. Having verified the robustness of CC700 for the purification of filamentous γPFD, we examined if the filaments could be directly functionalized in bacterial lysates and purified as complete nanostructures using CC700 resin. Previously, γPFD has been decorated with varying amounts of the fluorescent proteins mCerulean3 (CFP) and mVenus (YFP) via SpyTag/SpyCatcher conjugation to demonstrate the controlled positioning of functional domains (1). When aligned on γPFD filaments in nanometer-scale distances (<10 nm), the energy from an excited CFP is transferred to YFP through Förster Resonance Energy Transfer (FRET), resulting in decreased CFP emission and increased YFP emission. Varying the ratio of the two fluorescent proteins relative to the γPFD subunits enabled a controlled distribution and spacing of the fluorescent proteins along the filament. In this approach, three fusion proteins are recombinantly expressed and individually purified, including: γPFD C-terminally fused to a SpyTag domain (γPFD-SpyT), CFP C-terminally fused to a SpyCatcher domain (CFP-SpyC), and YFP C-terminally fused to a SpyCatcher domain (YFP-SpyC). Functionalized filaments were created by first assembling γPFD-SpyT into filaments, before mixing with CFP-SpyC and YFP-SpyC that conjugate to the SpyTags protruding from the γPFD-SpyT domains. However, this approach to build functionalized nanostructures was laborious and required each protein subunit to be separately purified and protein concentration determined to ensure appropriate molar ratios of the different protein subunits are combined. Furthermore, excess CFP-SpyC or YFP-SpyC that was not conjugated to γPFD-SpyT or excess γPFD-SpyT that was not functionalized was present in some mixtures, which requires additional purification steps for their removal.

In contrast to the previously published method, using CC700 should enable the purification of the functionalized filaments and the removal of unincorporated CFP and YFP subunits in the same step. We therefore adapted the SpyTag/SpyCatcher conjugation strategy for the functionalization of γPFD in bacterial lysates. In addition to the γPFD-SpyT, CFP-SpyC, and YFP-SpyC fusion proteins, a fourth protein will be recombinantly produced, the SpyCatcher domain (SpyC) that will be used to control the spacing of fluorescent subunits on the filament (Figure 3). In this new approach, filament functionalization will be achieved by mixing bacterial lysates that contain recombinant γPFD-SpyT filaments with lysates each containing SpyC, CFP-SpyC, and YFP-SpyC, followed by purification of functionalized filaments by MMC. Varying the relative amounts of each lysate should modulate the average distance between the two fluorescent proteins aligned on filaments, and thereby modulate the FRET signal produced by functionalized filaments (Figure 4a). Control over the average distance will be achieved by using the SpyC component to block SpyTags protruding from the filamentous γPFD-SpyT and thereby render these domains unavailable to bind the SpyCatcher domains of either CFP-SpyC or YFP-SpyC (Figure 3). Therefore, the amount of FRET will be inversely proportional to the amount of SpyC lysate added to the mixture of γPFD-SpyT, CFP-SpyC, and YFP-SpyC lysates. The use of the additional SpyC component to block SpyTags has the additional advantage of enabling assembly conditions to contain excess concentrations of CFP-SpyC and YFP-SpyC to ensure near complete γPFD functionalization.

**Figure 3.**
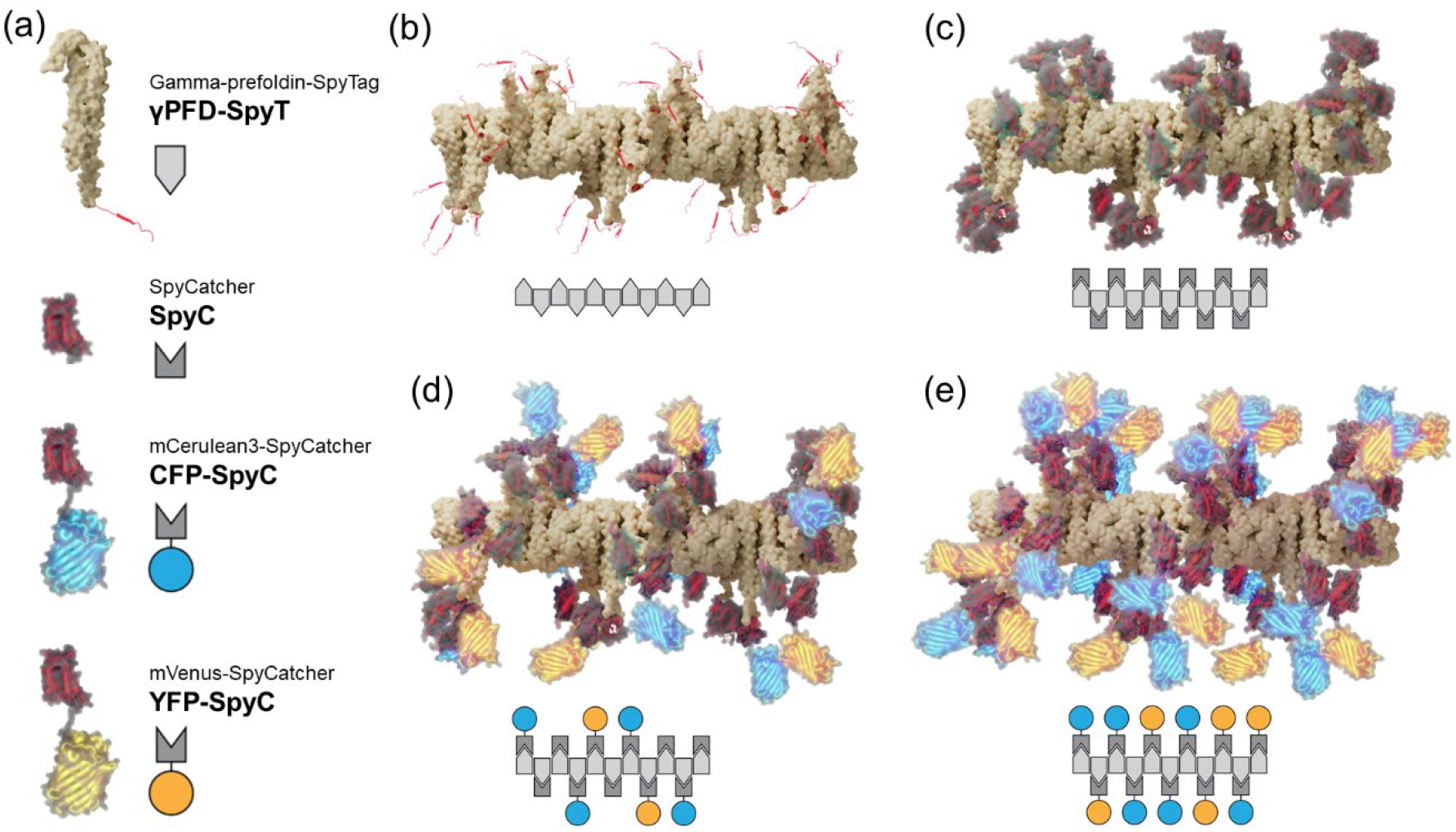
A strategy to attach and interspace functional proteins to γPFD filaments. **(a)** Predicted structural protein models of γPFD-SpyT and the functionalization subunits that are also represented as schematic icons. **(b)** Structural model of a γPFD-SpyT filament with SpyTag domains protruding from the C-termini of each subunit. **(c)** The SpyTag domains on γPFD-SpyT filaments can be blocked by the covalent attachment of SpyC subunits. **(d)** Attachment of CFP-SpyC and YFP-SpyC to the γPFD-SpyT filament via SpyTag/SpyCatcher covalent interactions. In this model, the CFP-SpyC and YFP-SpyC are randomly interspaced by SpyC. **(e)** Maximal density of CFP-SpyC and YFP-SpyC subunits attached to a γPFD-SpyT filament should be achieved by omitting the SpyC subunit.

**Figure 4.**
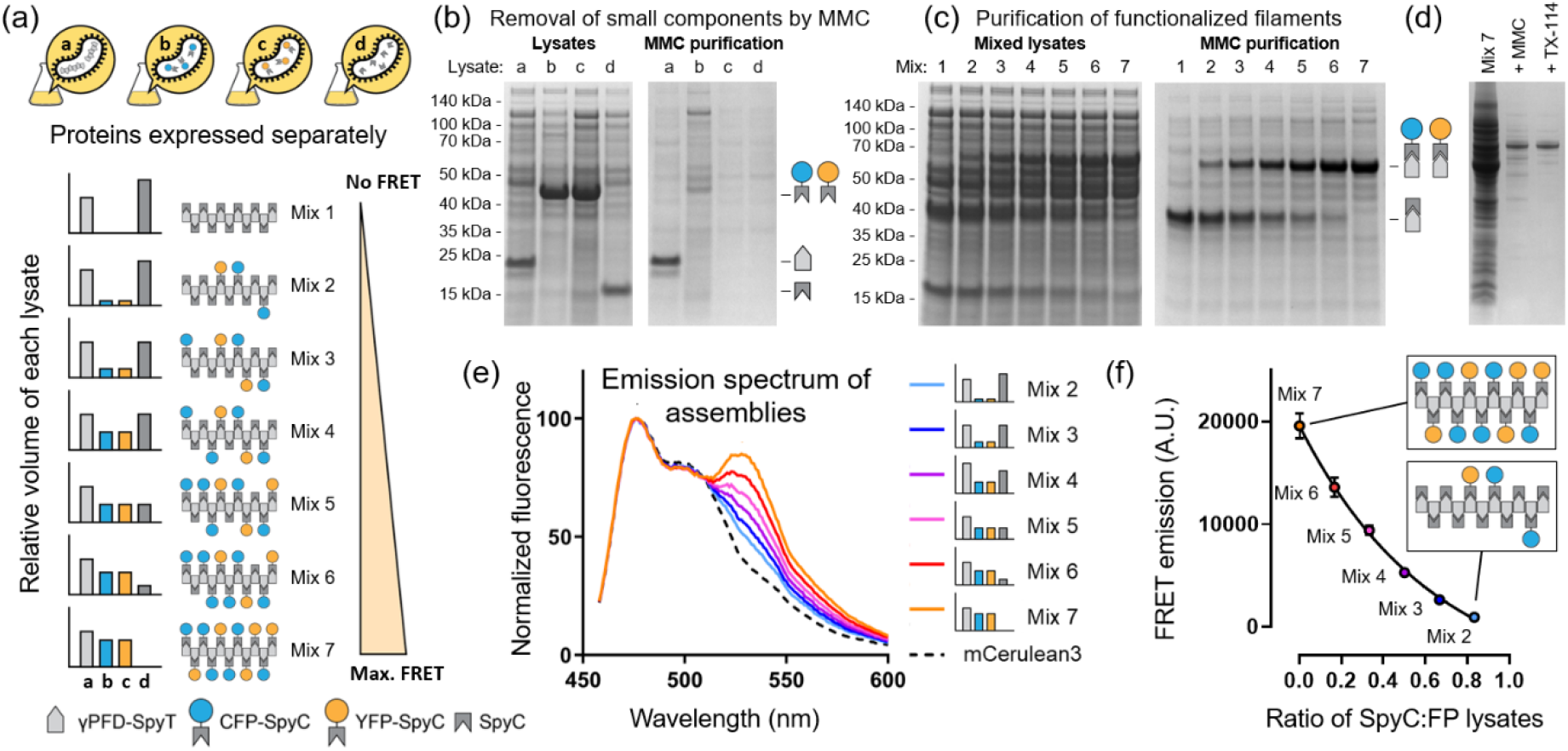
In vitro assembly and MMC purification of a functionalized γPFD protein filament. **(a)** Strategy for the assembly of functionalized protein filaments in bacterial lysates. The bars indicate the relative volumes of each lysate used in the different mixtures. Increasing the ratio of SpyC in the lysate mixture interspaces the fluorescent proteins CFP-SpyC and YFP-SpyC attached to γPFD-SpyT. **(b)** SDS-PAGE showing recombinant protein in bacterial lysates and after MMC. The large γPFD-SpyT filaments (lane a) pass through the resin whereas the small protein components are retained in the resin. **(c)** Lysate mixtures before and after MMC. Lysate mixtures demonstrate the conjugation of varying ratios of SpyC and the fluorescent proteins CFP-SpyC and YFP-SpyC to filaments and subsequent purification using MMC. **(d)** Improved purification of functionalized filaments using a larger volume of CC700 resin followed by TX-114 phase separation. **(e)** Fluorescence emission spectra of each functionalized filament preparation. Decreasing the amount of SpyC increases the likelihood that CFP-SpyC and YFP-SpyC are located close to each other on the γPFD-SpyT filament, increasing FRET (indicated by the higher emission at 527 nm relative to 476 nm). **(f)** The FRET emission of each filament preparation exhibits a strong correlation with the amount of SpyT used as a spacer between the fluorescent proteins. Data points represent the average of two experiments.

### Single-step purification of a functionalized protein filament assembled *in vitro*

DNA plasmids were prepared encoding the recombinant expression of the four recombinant proteins γPFD-SpyT, CFP-SpyC, YFP-SpyC, and SpyC. The successful expression of the four proteins in *E. coli* was confirmed by SDS-PAGE of the lysates (Figure 4b). Each lysate was subjected to MMC which showed that, as expected, the megadalton γPFD-SpyT filaments could be purified using CC700 resin. However, the CFP-SpyC, YFP-SpyC, and SpyC proteins were retained in the CC700 resin (Figure 4b). These results confirmed that in addition to removing bacterial proteins, CC700 can also be used to remove unincorporated small protein components of a protein nanostructure.

Next, γPFD filaments were functionalized with fluorescent proteins by mixing the four different bacterial lysates in varying volumetric ratios to demonstrate control over the spacing between functional domains. For these mixtures, the volume of lysate containing γPFD-SpyT was kept constant and the volumes of lysates containing CFP-SpyC, YFP-SpyC, and SpyC were varied. From a qualitative analysis of the SDS-PAGE results (Figure 4b), we observed that the CFP-SpyC, YFP-SpyC, and SpyC functional components were expressed in equal or higher amounts than γPFD-SpyT. Lysate volumes were chosen that ensured an excess of the functional components relative to γPFD-SpyT, with the relative volumes of lysates used for each mixture being illustrated in Figure 4a.

Following the mixing and incubation of lysates, conjugation between γPFD-SpyT and the other SpyCatcher-containing components was verified by SDS-PAGE (Figure 4c). An SDS-PAGE gel showed the formation of higher molecular weight species corresponding to a covalent attachment between γPFD-SpyT subunits of filaments and SpyC, CFP-SpyC or YFP-SpyC to form, respectively, γPFD-SpyT/C, γPFD-Spy-CFP and γPFD-Spy-YFP. Importantly, bands corresponding to excess SpyC, CFP-SpyC and YFP-SpyC were observed by SDS-PAGE, which are components not attached to γPFD-SpyT and therefore not incorporated into the protein filaments. Moreover, the band corresponding to γPFD-SpyT was not observed in the lysate mixtures, confirming that γPFD-SpyT subunits were completed conjugated with either of the three other subunits, and therefore the limiting component in the functionalization reactions.

The presence of unincorporated components in the lysates enables evaluation of the ability of CC700 to remove unincorporated components of a protein nanostructure. The lysates mixtures containing the protein assemblies were subjected to MMC purification (Figure 4c). The γPFD-SpyT/C, γPFD-Spy-CFP and γPFD-Spy-YFP subunits from the different assemblies were able to be isolated by MMC, whereas the excess SpyC, CFP-SpyC and YFP-SpyC were almost completely retained in the MMC beads. The MMC purification results suggested that functionalized filaments were created upon mixing of the lysates, that the filaments were composed of fluorescent (γPFD-Spy-CFP and γPFD-Spy-YFP) and non-fluorescent (γPFD-SpyT/C) conjugated subunits, and that the composition of the functionalized filaments were proportionate to the volumetric ratios of the different lysates used for functionalization.

The MMC purification results (Figure 4c) showed a limited amount of background contamination. A second round of MMC demonstrated the removal of these bacterial protein contaminates, and the γPFD-SpyT functionalized with only CFP-SpyC and YFP-SpyC was further purified by TritonX-114 phase separation to remove lipidic vesicles from the MMC flow through (Figure 4d). The morphology of the CFP-SpyC and YFP-SpyC functionalized γPFD-SpyT filaments was examined using by TEM (Figure S6). Filaments of γPFD-SpyT that were functionalized with the fluorescent proteins differed in morphology to non-functionalized filaments, which resembled concatenated “beads”. A FRET assay was used to confirm co-purification of the fluorescent proteins and non-fluorescent SpyCatcher protein with the filaments was due to their incorporation into filament nanostructures. FRET should occur when the mCerulean3 of γPFD-Spy-CFP subunits is less than 10 nm from the mVenus of γPFD-Spy-YFP. The likelihood of the donor and acceptor fluorescent subunits aligned within 10 nm along the filament scaffold should decrease as the number of the non-fluorescent γPFD-SpyT/C subunits increase, and the fluorescent subunits become progressively interspaced. Therefore, the lysate mixtures containing more CFP-SpyC and YFP-SpyC relative to SpyC should result in filaments that produce more FRET signal (Figure 4a). A 433 nm light source was used to excite CFP and the emission spectrum was recorded for each assembly (Figure 4e). The emission at 533 nm (optimal emission of YFP) relative to emission at 476 nm (optimal emission of CFP) increased with a composition higher in fluorescent subunits, which is indicative of increasing FRET. The deconvoluted FRET signal for each assembly is shown in Figure 4f and illustrates the steady decrease in FRET signal as the fluorescent subunits become increasingly interspaced by γPFD-SpyT/C subunits. In summary, the composition of the filaments could be controlled by combining varying volumes of bacterial lysates that contain functional domains and the resulting functionalized filaments purified in a single step using CC700. Using standard procedures, the functionalized γPFD assemblies would have required four separate purifications (and potentially buffer-exchanging steps) to produce the nanostructure components, followed by additional polishing steps to remove excess free components after assembly. Using CC700, no purification of the individual components was required, and purification of the large assembly was achieved concomitantly to the removal of unincorporated subunits in a single step. In our experiments, a second round of MMC was performed to improve purification, however the same result could be obtained through the use of a CC700 column with a greater volume of resin and binding capacity to increase the purity of nanostructures in eluted early fractions. Thus, the production of protein nanostructures can be greatly accelerated by use of CC700 resin as the method of purification.

## Conclusions

Herein, we have demonstrated a simple MMC method using CC700 to purify recombinantly produced protein nanostructures. Our method bypasses the need to separately purify each protein component, works in a wide range of biocompatible solvents, and removes both bacterial proteins and excess recombinant proteins that do not integrate the nanostructure in a single step. In the present study, these experiments have been performed at a small scale, however the method is inherently scalable. Furthermore, we presented both pre-processing (TFF) and post-processing (TX-114 phase separation) methods that can be used to improve the quality of the purification. Many more downstream polishing procedures exist (23) that should be compatible with CC700, especially in the context of large-scale purifications.

We also envision that with careful genetic encoding of the protein components, it may be possible to fully assemble complex protein nanostructures *in vivo* via the co-expression of the protein components, followed by purification of the nanostructures by MMC. Such a strategy could further reduce the time and costs to produce protein nanodevices. Orthogonal protein interfaces and Tag/Catcher domains have been shown to guide the association of proteins *in vivo* (11, 24–27), opening a path for the *in vivo* preparation of protein nanostructures and nanodevices. A major challenge will be the control of the total and relative expression levels of each protein component to ensure the correct self-assembly and minimize the excess of free protein components that do not partake in the assembly. Novel strategies for the regulation of protein co-expression (28, 29) will need to be explored to achieve this vision.

## Declarations

### Consent for publication

All authors consent to the publication of this manuscript.

### Competing interests

The authors declare no competing interests.

### Author contributions

DLW and HL designed and performed experimental procedures, except transmission electron microscopy. JBM performed transmission electron microscopy and created the structural models. All authors contributed to the preparation of the manuscript.

## Acknowledgements

Protein purification and transmission electron microscopy were performed at the Recombinant Products Facility (RPF) and Electron Microscope Unit (EMU), respectively, in the Mark Wainwright Analytical Centre.

## Funding

This work was supported by the Air Force Office of Scientific Research (FA9550-20-1-0389).

## Abbreviations

γPFD: gamma-prefoldin
CC700: Capto Core 700
CFP: Cyan fluorescent protein
FP: Fluorescent protein
MMC: multimodal chromatography
PPI: protein-protein interaction
SpyC: SpyCatcher
SpyT: SpyTag
TFF: tangential flow filtration
TX-114: Triton X-114
YFP: Yellow fluorescent protein

## Notes

### Competing Interest Statement

The authors have declared no competing interest.

## References

1. Lim S, Jung GA, Glover DJ, Clark DS. Enhanced Enzyme Activity through Scaffolding on Customizable Self-Assembling Protein Filaments. Small. 2019;15(20):e1805558.

2. Wei Q, He S, Qu J, Xia J. Synthetic Multienzyme Complexes Assembled on Virus-like Particles for Cascade Biosynthesis In Cellulo. Bioconjug Chem. 2020;31(10):2413–20.

3. Zhang G, Johnston T, Quin MB, Schmidt-Dannert C. Developing a Protein Scaffolding System for Rapid Enzyme Immobilization and Optimization of Enzyme Functions for Biocatalysis. ACS Synth Biol. 2019;8(8):1867–76.

4. Lemmens LJM, Ottmann C, Brunsveld L. Conjugated Protein Domains as Engineered Scaffold Proteins. Bioconjug Chem. 2020;31(6):1596–603.

5. Chen YX, Ing NL, Wang F, Xu D, Sloan NB, Lam NT, et al. Structural Determination of a Filamentous Chaperone to Fabricate Electronically Conductive Metalloprotein Nanowires. ACS Nano. 2020;14(6):6559–69.

6. Luo Q, Hou C, Bai Y, Wang R, Liu J. Protein Assembly: Versatile Approaches to Construct Highly Ordered Nanostructures. Chem Rev. 2016;116(22):13571–632.

7. Cannon KA, Ochoa JM, Yeates TO. High-symmetry protein assemblies: patterns and emerging applications. Curr Opin Struct Biol. 2019;55:77–84.

8. Kuan SL, Bergamini FRG, Weil T. Functional protein nanostructures: a chemical toolbox. Chem Soc Rev. 2018;47(24):9069–105.

9. Reinke AW, Grant RA, Keating AE. A synthetic coiled-coil interactome provides heterospecific modules for molecular engineering. J Am Chem Soc. 2010;132(17):6025–31.

10. Thompson KE, Bashor CJ, Lim WA, Keating AE. SYNZIP protein interaction toolbox: in vitro and in vivo specifications of heterospecific coiled-coil interaction domains. ACS Synth Biol. 2012;1(4):118–29.

11. Chen Z, Boyken SE, Jia M, Busch F, Flores-Solis D, Bick MJ, et al. Programmable design of orthogonal protein heterodimers. Nature. 2019;565(7737):106–11.

12. Veggiani G, Nakamura T, Brenner MD, Gayet RV, Yan J, Robinson CV, et al. Programmable polyproteams built using twin peptide superglues. Proc Natl Acad Sci U S A. 2016;113(5):1202–7.

13. Swartz AR, Chen W. SpyTag/SpyCatcher Functionalization of E2 Nanocages with Stimuli-Responsive Z-ELP Affinity Domains for Tunable Monoclonal Antibody Binding and Precipitation Properties. Bioconjug Chem. 2018;29(9):3113–20.

14. James KT, Cooney B, Agopsowicz K, Trevors MA, Mohamed A, Stoltz D, et al. Novel High-throughput Approach for Purification of Infectious Virions. Sci Rep. 2016;6:36826.

15. Lagoutte P, Mignon C, Donnat S, Stadthagen G, Mast J, Sodoyer R, et al. Scalable chromatography-based purification of virus-like particle carrier for epitope based influenza A vaccine produced in Escherichia coli. J Virol Methods. 2016;232:8–11.

16. Weigel T, Solomaier T, Peuker A, Pathapati T, Wolff MW, Reichl U. A flow-through chromatography process for influenza A and B virus purification. J Virol Methods. 2014;207:45–53.

17. Lagoutte P, Mignon C, Stadthagen G, Potisopon S, Donnat S, Mast J, et al. Simultaneous surface display and cargo loading of encapsulin nanocompartments and their use for rational vaccine design. Vaccine. 2018;36(25):3622–8.

18. Song Y, Rodgers VG, Schultz JS, Liao J. Protein interaction affinity determination by quantitative FRET technology. Biotechnol Bioeng. 2012;109(11):2875–83.

19. Holstein MA, Nikfetrat AA, Gage M, Hirsh AG, Cramer SM. Improving selectivity in multimodal chromatography using controlled pH gradient elution. J Chromatogr A. 2012;1233:152–5.

20. Glover DJ, Lim S, Xu D, Sloan NB, Zhang Y, Clark DS. Assembly of Multicomponent Protein Filaments Using Engineered Subunit Interfaces. ACS Synth Biol. 2018;7(10):2447–56.

21. Glover DJ, Giger L, Kim SS, Naik RR, Clark DS. Geometrical assembly of ultrastable protein templates for nanomaterials. Nat Commun. 2016;7:11771.

22. Taguchi Y, Schatzl HM. Small-scale Triton X-114 Extraction of Hydrophobic Proteins. Bio Protoc. 2014;4(11).

23. Zydney AL. Continuous downstream processing for high value biological products: A Review. Biotechnol Bioeng. 2016;113(3):465–75.

24. Bedbrook CN, Kato M, Ravindra Kumar S, Lakshmanan A, Nath RD, Sun F, et al. Genetically Encoded Spy Peptide Fusion System to Detect Plasma Membrane-Localized Proteins In Vivo. Chem Biol. 2015;22(8):1108–21.

25. Giessen TW, Silver PA. A Catalytic Nanoreactor Based on in Vivo Encapsulation of Multiple Enzymes in an Engineered Protein Nanocompartment. Chembiochem. 2016;17(20):1931–5.

26. Wang XW, Zhang WB. SpyTag-SpyCatcher Chemistry for Protein Bioconjugation In Vitro and Protein Topology Engineering In Vivo. Methods Mol Biol. 2019;2033:287–300.

27. Smith AJ, Thomas F, Shoemark D, Woolfson DN, Savery NJ. Guiding Biomolecular Interactions in Cells Using de Novo Protein-Protein Interfaces. ACS Synth Biol. 2019;8(6):1284–93.

28. Meyer AJ, Segall-Shapiro TH, Glassey E, Zhang J, Voigt CA. Escherichia coli “Marionette” strains with 12 highly optimized small-molecule sensors. Nat Chem Biol. 2019;15(2):196–204.

29. Rouches MV, Xu Y, Cortes LBG, Lambert G. A plasmid system with tunable copy number. Nat Commun. 2022;13(1):3908.

