## Supplemental data and figures for "Single-step purification of functionalized protein nanostructures using multimodal chromatography"

Supplemental data for "**Single-step purification of functionalized protein nanostructures using multimodal chromatography**"

Daniel L. Winter, Hélène Lebhar, Joshua B. McCluskey, and Dominic J. Glover

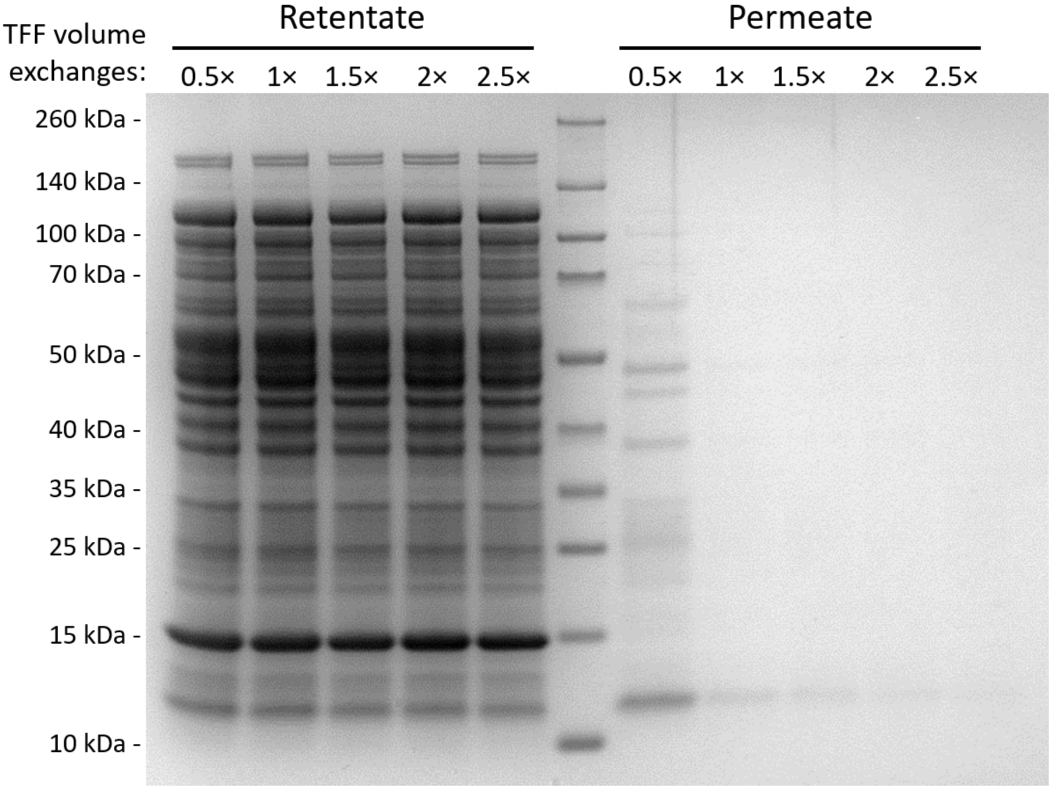


**Figure S1. SDS-PAGE analysis of the retentate and permeate fractions produced during tangential flow filtration**. A small quantity of bacterial proteins passes through the membrane during the early stages of TFF (less than one volume). By two volumes, the system has nearly reached equilibrium. The protein content in the retentate is virtually unchanged throughout the whole procedure.

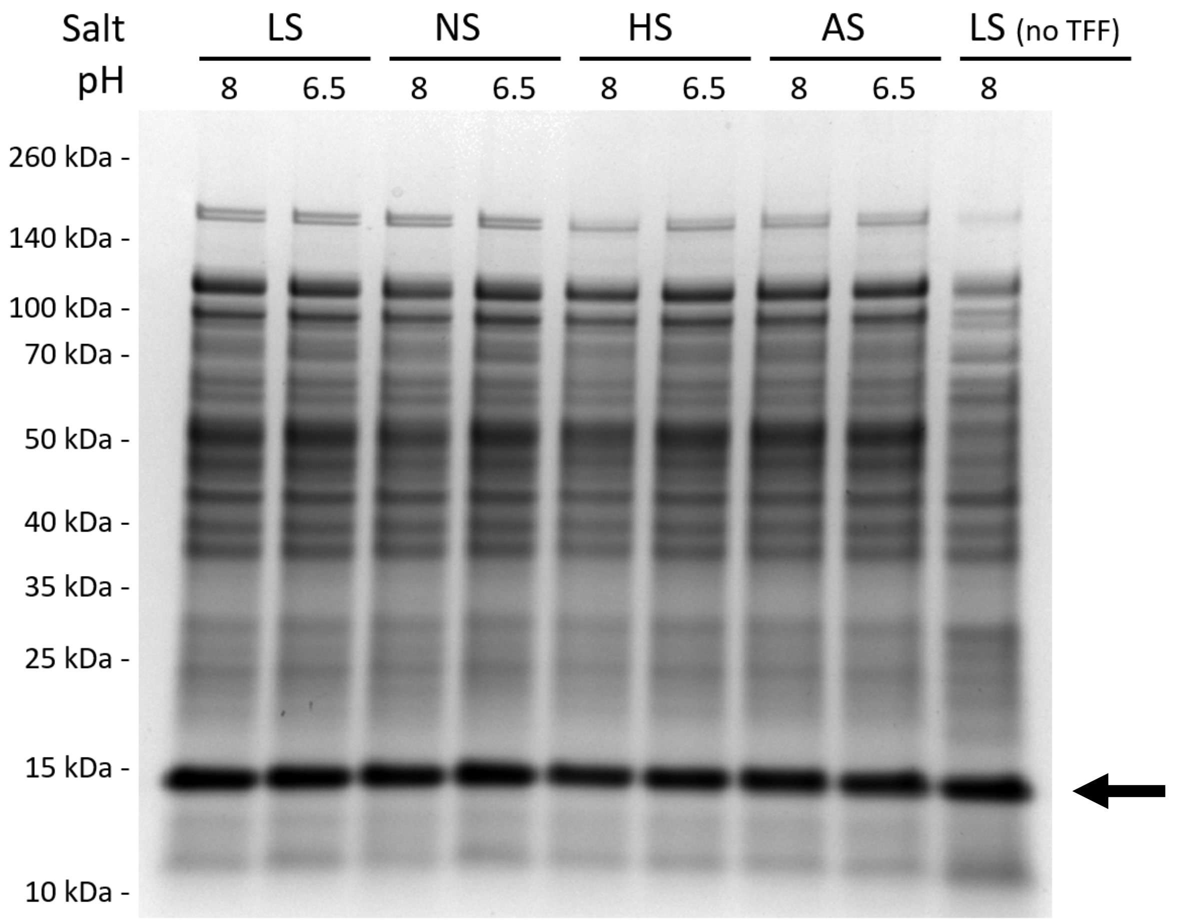


**Figure S2. Lysate samples used for multimodal chromatography purification of gamma-prefoldin.** Cell lysate was prepared by lysis of *E. coli* cells in a low salt buffer (pH 8), followed by DNase treatment and clarification by centrifugation (rightmost lane). Subsequently, the lysate was subjected to TFF (leftmost lane). The TFF-processed sample was then aliquoted and dialyzed into other buffers that vary in salt and pH conditions. None of these procedures significantly changed the total protein content, nor the amount of gamma-prefoldin (arrow), in each sample. LS: low salt; NS: no salt; HS: high salt; AS: ammonium sulphate. See Table 1 for more details on each buffer condition.

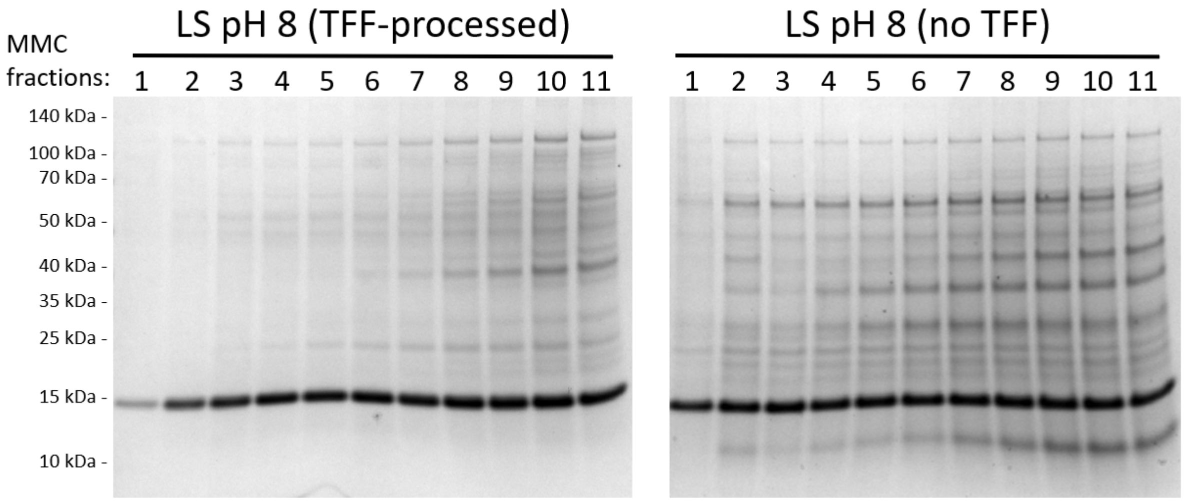


**Figure S3. Comparison of the early MMC elution fractions between samples that had been processed or not processed previously by tangential flow filtration.** The first eleven fractions (corresponding to 11 column volumes) from a MMC purification of recombinant gamma-prefoldin from bacterial lysate were analyzed by SDS-PAGE. TFF pre-processing (gel on the left) of the lysate resulted in higher purity and delayed saturation of the resin (indicated by the appearance of protein contaminants) compared to the same lysate without TFF pre-processing (gel on the right).

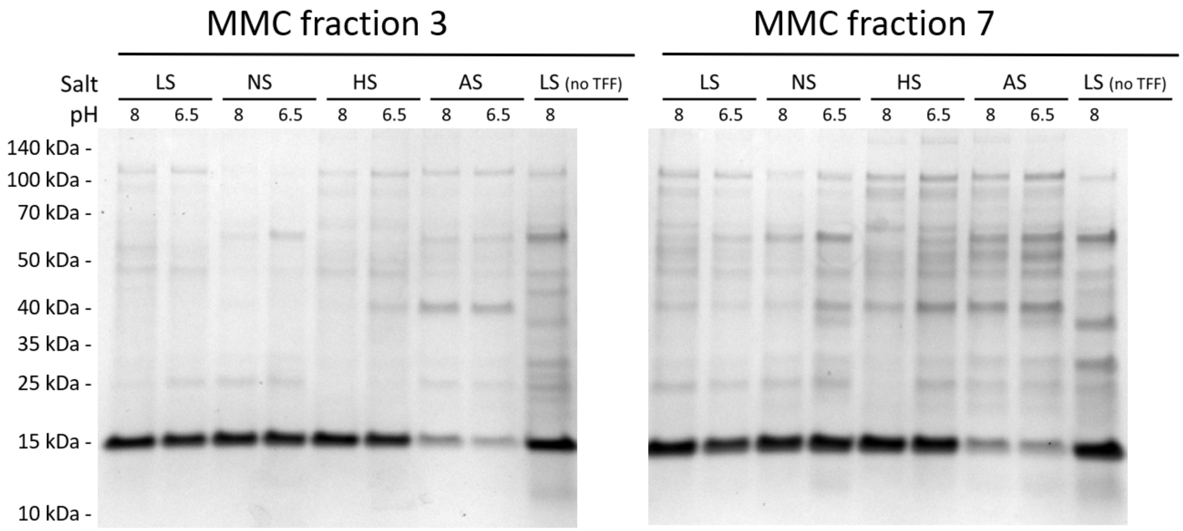


**Figure S4. Comparison of initial fractions from MMC purification of recombinant gamma-prefoldin performed with different buffered solvents.** The comparison of the third elution fraction from each MMC run (also shown in Figure 2d) shows that MMC performs comparably in all solvents at the early stages of MMC, that ammonium sulfate caused some precipitation of gamma-prefoldin, and that omitting TFF pre-processing of the lysate in lesser purity of gamma-prefoldin. A comparison of a later fraction (the seventh elution fraction) reveals slight differences in performance between solvents, with some fractions exhibiting the first signs of resin saturation (greater amounts of bacterial protein contamination).

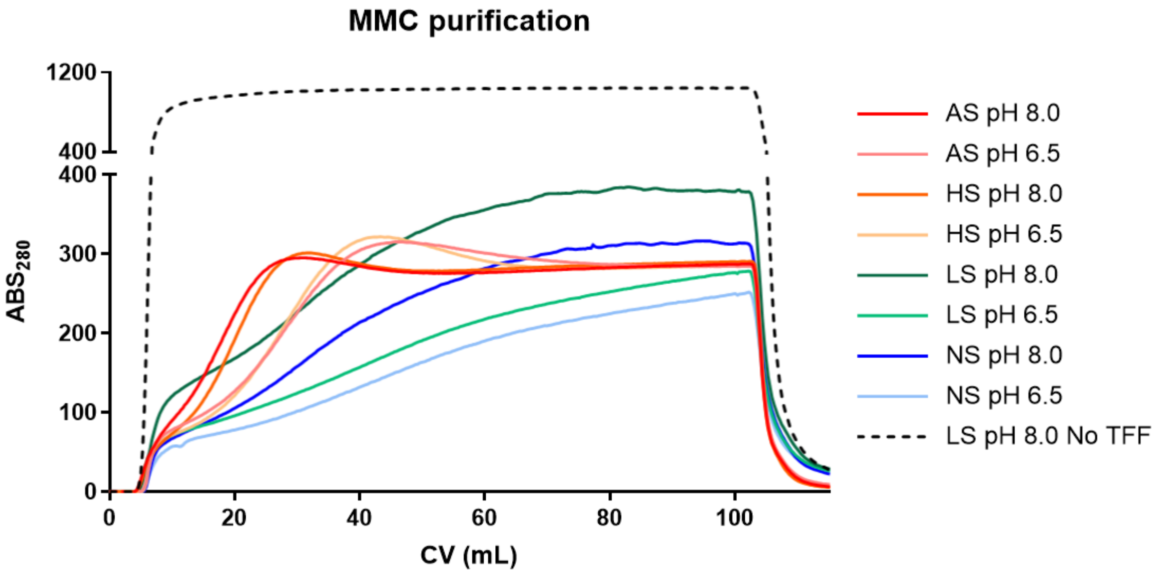


**Figure S5. Extended UV chromatograms of MMC purifications of recombinant gamma-prefoldin with varying buffer conditions.** The binding capacity of MMC for gamma-prefoldin is comparable in a range of buffered solvents during the early stages of chromatography. The onset of resin saturation is approximately 5 column volumes for all solvents (see Figure 2c for a detailed view). Omitting the TFF pre-processing (dashed line) results in an earlier and steeper onset of resin saturation.

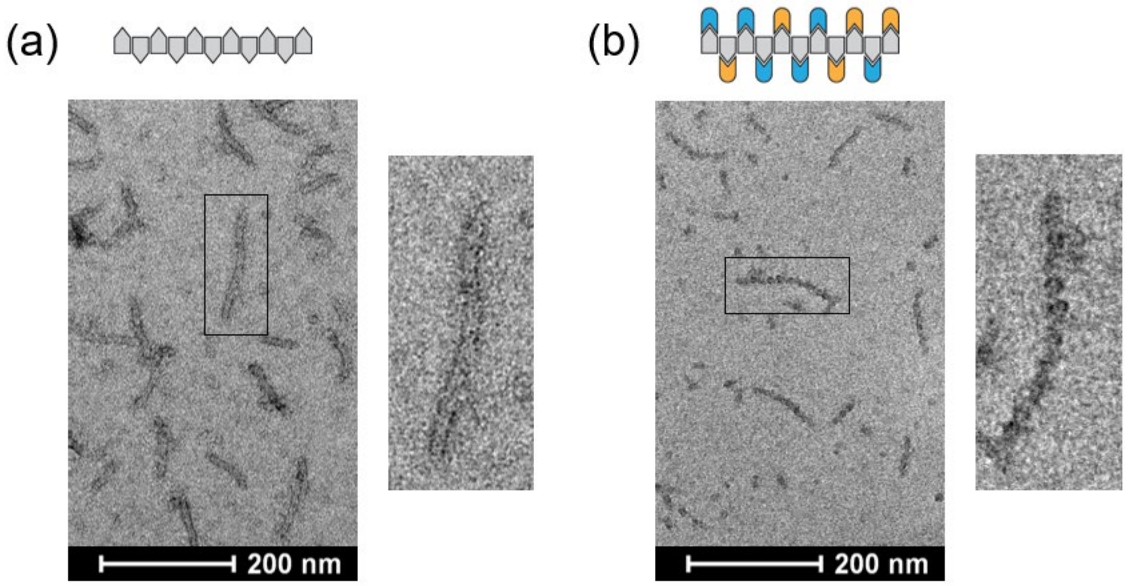


**Figure S6. Transmission electron microscopy of gamma-prefoldin shows differences in the morphology of non-functionalized and functionalized protein filaments.** (a) Gamma-prefoldin-SpyTag (γPFD-SpyT) forms filaments with similar morphology to wild type gamma-prefoldin (see Figure 1f), although with shorter average filament lengths. (b) γPFD-SpyT functionalized with mCerulean3-SpyCatcher (mC3-SpyC) and mVenus-SpyCatcher (mV-SpyC) shows a different morphology, resembling concatenated “beads”.
